# Cryopreservation and revival of Hawaiian stony corals via isochoric vitrification

**DOI:** 10.1101/2023.03.05.531199

**Authors:** Matthew J. Powell-Palm, E. Michael Henley, Anthony N. Consiglio, Claire Lager, Brooke Chang, Riley Perry, Kendall Fitzgerald, Jonathan Daly, Boris Rubinsky, Mary Hagedorn

**Affiliations:** Department of Mechanical Engineering, Texas A&M University, College Station, TX, USA; Department of Materials Science and Engineering, Texas A&M University, College Station, TX, USA; Smithsonian National Zoo and Conservation Biology Institute, Front Royal, VA, 22630, USA; Hawai□i Institute of Marine Biology, University of Hawai□i at Mānoa, Kāne□ohe, HI, 96744, USA; Department of Mechanical Engineering, University of California Berkeley, Berkeley, CA, USA; Taronga Institute of Science and Learning, Taronga Conservation Society Australia, Mosman, NSW 2088, Australia; Centre for Ecosystem Science and Centre for Marine Science and Innovation, School of Biological, Earth and Environmental Sciences, University of New South Wales, Sydney, NSW 2052, Australia

## Abstract

Corals are under siege by both local and global threats, creating a wordwide reef crisis. Cryopreservation is an important intervention measure and a vital component of the modern coral conservation toolkit, but preservation techniques are currently limited to sensitive reproductive materials that can only be obtained a few nights per year during spawning. Here, we report the first successful cryopreservation and revival of cm-scale coral fragments via mL-scale isochoric vitrification. We demonstrate coral viability at 24h post-thaw using a calibrated oxygen-uptake respirometry technique, and further show that the method can be applied in a passive, electronics-free configuration. Finally, we detail a complete prototype coral cryopreservation pipeline, which provides a platform for essential next steps in modulating postthaw stress and initiating long-term growth. These findings pave the way towards an approach that can be rapidly deployed around the world to secure the biological genetic diversity of our vanishing coral reefs.

## Introduction

Coral reefs around the world are imperiled by local and global threats, leading to shocking declines in species and genetic diversity^1^. Over the last five decades, coral reef losses have accelerated^2^, and future losses are predicted at 70–99%^3^. Coral reefs are some of the oldest, most diverse ecosystems on our planet and act as an oceanwide nursery for a quarter of all marine life^4^. They protect coastlines, cities and homes, are critical to food security, and support widespread recreational activities^5,6^. Today, the economic value of the world’s coral reefs is estimated at $10 trillion USD per year^7^. However, despite their unparalleled global economic and social value, coral reefs continue to decline, threatening most life on Earth, and more imminently the lives and livelihoods of resource-poor but reef-rich nations^8^.

The coming decade will be critical for the future of coral reefs, and therefore broad scientific interventions are sought^9^. One such intervention action is cryopreservation. Cryopreserved coral sperm has been being banked for more than a decade and has been used to assist gene flow both in the critically endangered Caribbean Elkhorn coral^10^ and on the Great Barrier Reef^11^, and recent advances in novel laser rewarming techniques have also enabled cryopreservation of coral larvae^12^. However, while preserved reproductive materials can be utilized for reef restoration and diversification, climate change and its related bleaching events are negatively impacting coral reproduction^13–15^. Therefore, a climate change-resistant approach is needed to enable cryopreservation of the entire coral organism without waiting for increasingly uncertain yearly reproductive events. Towards that goal, we describe herein the first successful cryopreservation of whole cm-scale coral microfragments using a cutting-edge thermodynamic cryopreservation process called isochoric vitrification.

Scaling cryopreservation protocols from isolated cell suspensions (such as sperm) to large tissues or whole organisms presents a number of physical challenges, chief among them that these more complex systems (unlike their cell suspension counterparts) can generally tolerate little-to-no ice formation in the system. This therefore requires that they be cryopreserved in an ice-free “glassy” state, by the complex kinetic process of vitrification^16^. Vitrification is highly dependent on both system chemistry and cooling/warming rates^16,17^, and as sample size increases, the decreasing surface area-to-volume ratio sharply limits both surface-driven heat transport and the osmotic action of chemical cryoprotectants.

Recent work has provided a wellspring of new approaches that grapple with these scaling issues: optimized metallic meshes to enable ultrafast cooling rates in microscale samples, nanoparticle-enhanced optical and electromagnetic warming techniques to enable ultrafast and uniform warming, advances in perfusion technologies to enable loading of higher-osmolality, higher-viscosity cryoprotective solutions, etc.^18–21^. However, the many exciting solutions that have emerged in the past five years are unified by two common limitations: sample size (≪1mL) and/or tolerance of these samples to the highly concentrated (7-10M) and potentially toxic solutions required.

In this work, we present the first biological validation of a novel thermodynamic technique, isochoric vitrification^22^, that facilitates the vitrification process at low cooling and warming rates (<100°C/min and <400°C/min respectively) in bulk-volume samples (5 mL) of a relatively low-molarity (~4M) vitrification solution. We combine this technique with key advances in coral husbandry and evaluation to achieve cryopreservation and revival of the endemic Hawaiian coral *Porites compressa*, providing the first-ever demonstration of cryopreservation of mature coral fragments of ~1cm length scales. After vitrification, we demonstrate a 24h post-thaw survival of coral microfragments using oxygen-uptake respirometry. Finally, we show that this technique can be successfully applied without active monitoring, in an electronics-free isochoric chamber ready for use in the field. We suggest that this technique can be applied any time of year and in most relevant environments and thus holds great promise for ensuring the maintenance of coral diversity.

## Results

### Isochoric vitrification: thermodynamic premise and validation

Vitrification techniques typically employ isobaric (constant-pressure) thermodynamic conditions, wherein the system (i.e., the biological sample of interest and surrounding preservation media) is open to the atmosphere or another reservoir of pressure. Under these conditions, the uniquely large density difference between liquid water and ice-Ih does not appreciably affect the kinetic processes of ice nucleation and vitrification. However, if the system is rigidly confined and *denied* access to the atmospheric pressure reservoir, i.e., held under isochoric (constant-volume) conditions, both the equilibrium thermodynamics and the kinetics of nucleation and growth change dramatically^22–27^. Under confinement, the expansion exacts a sizable energetic toll on the nascent ice phase, which has been shown in theory and experiment to both limit the ultimate growth of ice^23,24^ and reduce the probability of its initial nucleation from arbitrary supercooled aqueous systems^25–27^. Avoiding the formation of ice at temperatures between the melting point and the glass transition is the central challenge in facilitating vitrification, and we thus hypothesized that the same suite of physical effects driving ice avoidance in the past decade of high subzero isochoric experiments may also help to facilitate vitrification in isochoric systems^22^.

In order to probe this supposition, we designed a custom isochoric vitrification chamber fitted with a pressure transducer (Fig. 1a) and leveraged the principle of pressure-based isochoric nucleation detection^26^ to evaluate the isochoric vitrification behaviors of a new coral preservation solution. We based this solution, called CVS1 (1.05M DMSO, 1.05M Glycerol, 1.05M PG, 0.85M trehalose in phosphate buffered saline; total molarity 4.15; more details in SI), on a solution used previously to preserve coral larvae at microliter volumes with the assistance of laser rewarming^12^. While the bulk vitrification process is typically evaluated visually^28^, the pressure generated by ice formation in isochoric systems provides a more quantitative tool by which to evaluate vitrification protocols, with both the presence of a pressure spike and its magnitude providing valuable physical information.

**Figure 1.**
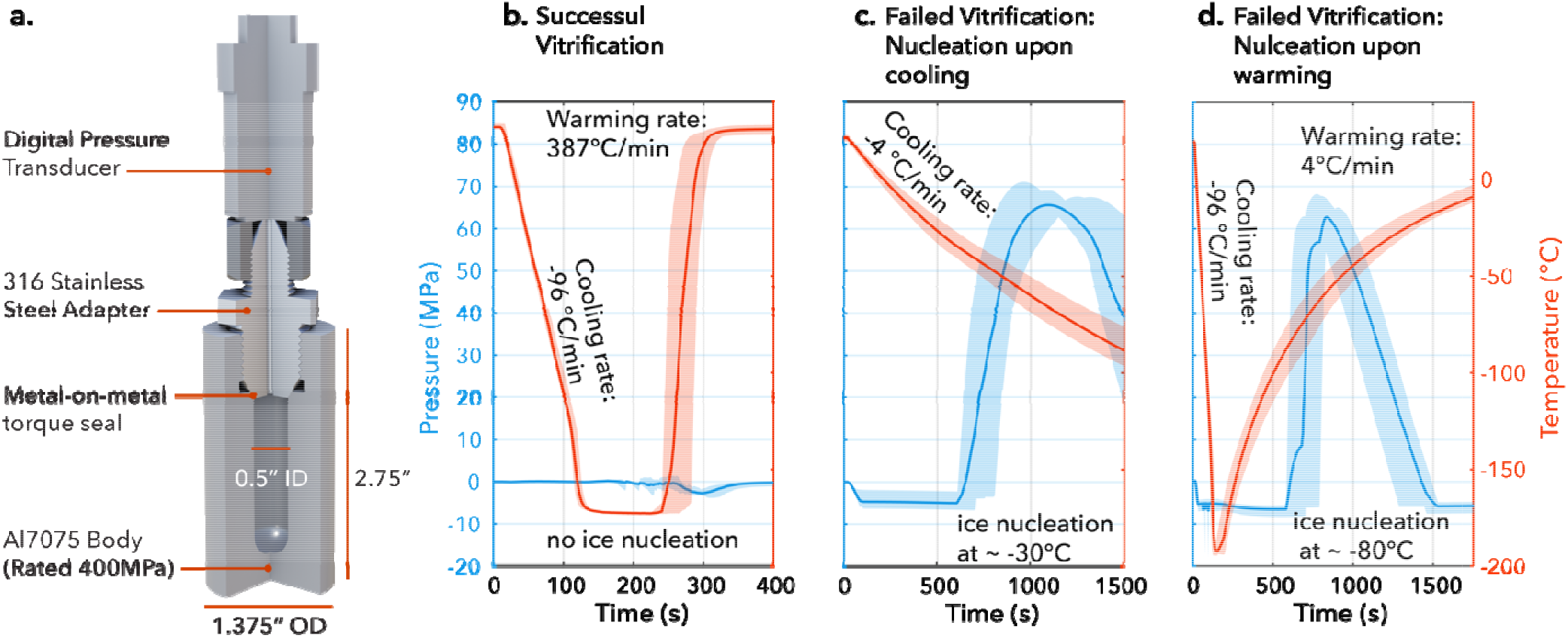
Facilitating and verifying an isochoric vitrification process. a) Custom-designed isochoric chamber rated for 400 MPa / −200 °C. The chamber body is built of aluminum 7075-T651 for its combination of excellent thermal and mechanical properties, and the threaded adapter is built of 316 stainless steel. The adapter possesses a significantly lower temperature coefficient of thermal expansion compared to Al-7075, thus producing an advantageous thermal shrink-seal upon cooling. b) Pressure-Temperature-Time (PTT) trace for successful vitrification, as indicated by the absence of a jump in pressure (n=10). c, d) PTT traces for slower cooling (c) and warming (d) rates, each of which produce ice nucleation and the according pressurization (n=10, each).

Fig. 1 demonstrates the thermodynamic evaluation process used to determine whether vitrification has occurred within the sealed chamber. In brief, the isochoric chamber, shown in Fig. 1a, is plunged into a bath of liquid nitrogen (up to the connection mating it to the pressure transducer, which is itself rigorously insulated to keep the encased electronics above −50 °C at all times), allowed to reach steady state, then plunged into a water bath at 27 °C with submersible pumps providing turbulent mixing. The temperature and pressure are monitored throughout, with a rapid increase in pressure indicating growth of ice in the system^23,26^. This simple plunge method provided maximum cooling/warming rates (i.e., at the periphery of the liquid sample) of 96 +/-6 °C/min and 387 +/- 55 °C/min respectively and did not produce the sharp increase in pressure indicative of ice nucleation (Fig 1b). CVS1 was thus concluded to vitrify successfully under these conditions. In order to verify this conclusion and explore the sensitivity of the vitrification process, we also performed the same experiments at slower cooling and warming rates, achieved by plunging the chamber with varying degrees of insulation. As shown in Fig 1c, reduced cooling rates elicited ice-driven pressurization upon cooling to approximately 65 MPa. Similarly, when the chamber was once again cooled at 96 °C/min but rewarmed more slowly, the same pressurization was observed (Fig. 1d). This cooling/warming rate dependence is the hallmark of vitrification– a kinetic process that depends sharply on the thermal history of the sample.

It should also be noted even at the mild cooling rates that do produce detectable ice growth, some degree of partial vitrification is likely occurring within the system, driven by the unique combination of pressure effects and solute ripening that is characteristic to the isochoric freezing process^29^.This is suggested by the local maximum in pressure within the plausible temperature vicinity of the glass transition, which implies a ceasing of ice growth and a gradual transition to the more contractile glassy state^30–32^ (See SI for supporting calculation and further discussion). The notion of isochoric partial vitrification thus introduces the intriguing premise that a biological sample within the system could potentially still vitrify even while faced with ice growth elsewhere in the chamber. However, this same competition between icy expansion and glassy contraction, and the additional effects of contraction of the chamber, place limits on the absolute sensitivity of pressure-based vitrification detection, e.g., even in the case of Fig. 1b wherein no net pressure increase is observed, some small amount of ice could still be forming. The intricacies of these thermo-volumetric effects should be explored in future work.

Finally, in order to determine whether the observed vitrification of the CVS1 solution is indeed a product of the employed isochoric thermodynamic conditions, as hypothesized, we repeated the trials shown in Fig. 1b with an unsealed chamber open to atmospheric pressure. The solution froze in all cases (indicated by significant expansion of the sample, as opposed to the significant contraction that accompanies vitrification^31^), confirming that isochoric confinement facilitates the observed vitrification process. Additional detailed description of all thermodynamic tests is included in the SI.

### Cryopreservation of Hawaiian stony coral fragments

Having validated that the CVS1 coral preservation solution vitrified at bulk volumes under isochoric conditions, we proceeded to test this process on the Hawaiian coral *Porites compressa.* Our experimental workflow is shown in Fig. 2, and described in further detail below, in the Methods, and in the SI.

**Figure 2.**
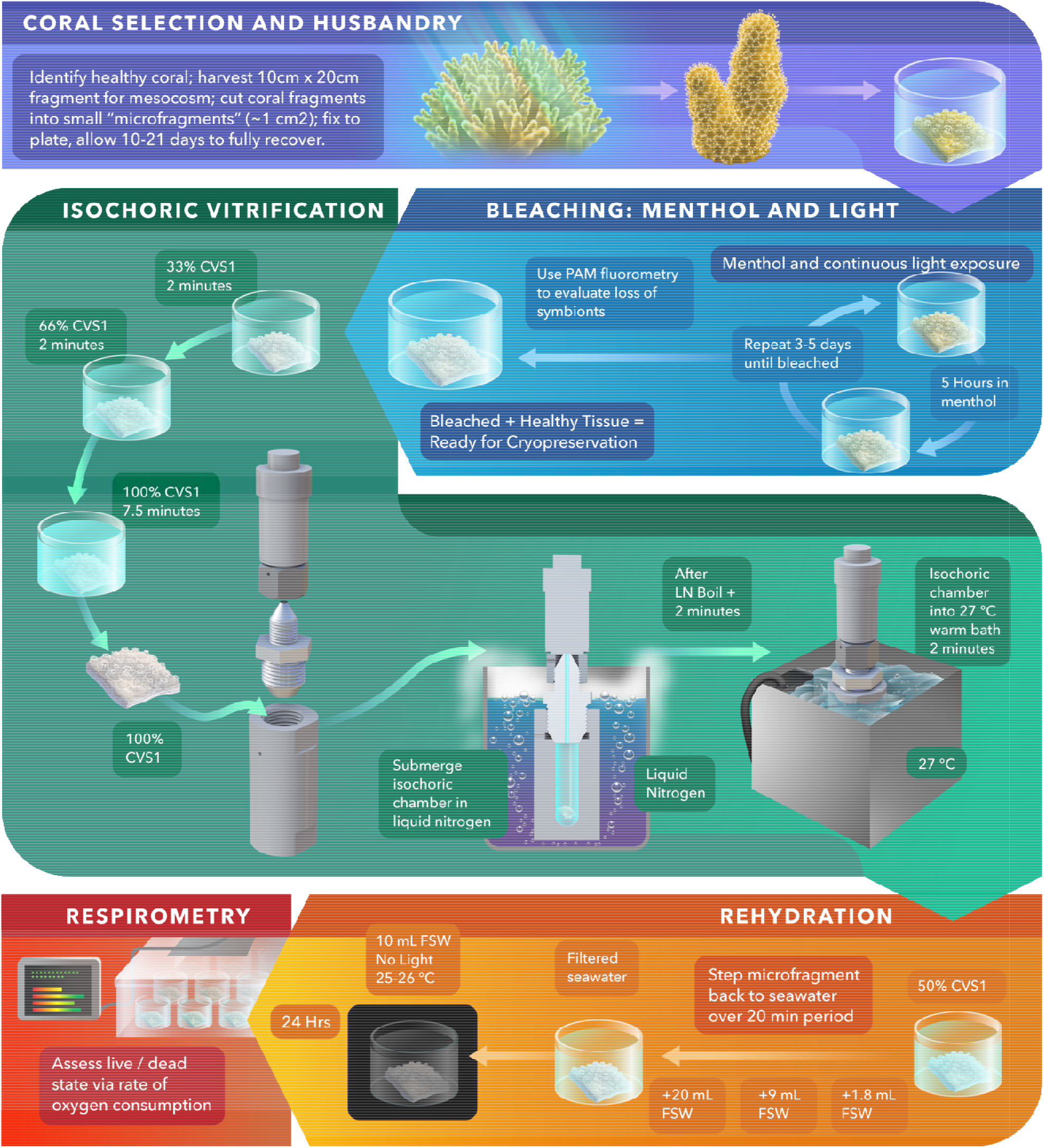
Coral cryopreservation and revival protocol. Each step shown here is described in brief in the text, and step-by-step procedures details are provided in the SI.

### Selection and fragmentation

First, a healthy adult coral colony is located on the reef, from which a fragment approximately 10cm x 20cm is collected and maintained in an outdoor flow-through mesocosm system at the Hawai□i Institute of Marine Biology located in Kāne□ohe Bay, Hawai□i. This fragment is then further cut into several ~1cm^2^ “microfragments”, each of which contains approximately 20 individual polyps, and glued to a gridded acrylic plate organized by genotype identification and date of microfragment processing. After a period of at minimum 10 and at maximum 30 days, the microfragments recover fully from the stress of fragmentation and are ready for use in the cryopreservation process.

### Bleaching

The microfragments are brought into the laboratory and then bleached with a combination of menthol and light, according to the protocol of Lager et al^33^ (detailed also in the SI). This causes them to shed their algal symbionts, which in previous work and preliminary trials were found to interfere with the CPA-penetration / coral tissue dehydration process due to the differences in cryobiological parameters between the symbionts and the tissue^34,35^. Sufficient absence of symbionts post-bleaching is confirmed by pulse amplitude modulation (PAM) fluorometry (see SI for full photosynthetic yield criteria).

### CPA loading

Bleached fragments (as shown in Fig. 2) are then introduced into full-concentration CVS1 in three submersion steps (33% CVS1, 2min; 66% CVS1, 2min, 100% CVS1, 3 – 7.5 min) in order to avoid significant osmotic stress. The composition of CVS1 was engineered to produce a combination of dehydration (driven by trehalose, which cannot penetrate the coral tissue) and CPA penetration (driven by DMSO and PG, which can both cross the tissue membrane). While the ultimate degrees of penetration and dehydration achieved cannot be quantified here, we hypothesize that dehydration is the dominant process, as coral tissue contains extremely fast-acting water channels, characterized by their low membrane permeability^12,35^. Furthermore, in order to probe the sensitivity of the ensuing coral vitrification process to successful dehydration/penetration from CVS1 (and to screen for any acute toxicity effects), we introduce the diffusion time in the full-strength solution as an experimental variable, testing exposure times of 3, 5, and 7.5 minutes.

### Isochoric vitrification of coral microfragments

Following CPA loading, a given coral microfragment is then placed into an isochoric chamber filled with CVS1 solution, sealed to a torque of 45 ft-lb, and promptly plunged into a bath of liquid nitrogen (LN_2_). This process requires approximately 30 seconds, and care is taken to ensure that no bulk air bubbles are introduced into the chamber during loading, as the presence of bulk air has been shown to both alter the isochoric thermodynamic path^36^ and aid in the nucleation of ice^37^. In our core suite of vitrification trials, the chamber was instrumented with a digital pressure transducer (Ellison Sensors Inc., USA) as shown in Fig 1a, and the exact protocol used to conduct the previous thermodynamic trials was performed, again employing the absence of an increase in pressure to confirm successful vitrification. The chamber is held in LN_2_ until steady state is achieved (indicated by the cessation of LN_2_ boiling). Two minutes thereafter, the chamber is transferred directly from the LN_2_ bath to a circulating water bath at 27 +/-1°C, where it is submerged for two additional minutes. This temperature represents the maximum at which *Porites compressa* will not experience additional heat-based stress, thus eliminating the possibility of overheating. After warming is complete, the chamber is unsealed, and the coral is transferred into a petri dish of 50% CVS1 in filtered seawater. This dish is further diluted with filtered seawater (FSW) in several discreet steps (see SI for full details) over the next 19.5 minutes, at which point the coral is transferred to a fresh dish of 10mL FSW and allowed to recover in darkness at 25-26°C for the next 24 hours.

### Respirometry Technique for Analyzing Coral Survival

Following experimental treatment, a quantitative technique by which to evaluate the health of coral fragments in real time was needed. While existing techniques are adequate in many circumstances, they fall short when external stressors (including thermal and chemical stressors) may elicit visible physiological responses in the coral that do not necessarily equate to cell death, tissue damage, or other long term deleterious effects. Our lab established the utilities and limits of some of these methods for *Porites compressa* in previous work^33^, concluding ultimately that simple visual morphological evaluation, brightfield microscopic evaluation, and even confocal fluorescent evaluation provided limited use in establishing the relative health of stressed coral fragments for these experiments. In need of a more robust and quantitative method by which to evaluate the health of coral fragments, we used a microrespirometry technique to record oxygen consumption rates over time in order to differentiate between healthy and severely injured coral or dead coral^38^.

Importantly, coral is considered a holobiont, comprised of coral tissues, algal symbionts (Symbiodiniaceae, which are removed by bleaching prior to vitrification), and a bacterial microbiome. We demonstrate here that the oxygen consumption balance between the bleached coral tissue and its bacterial microbiome provides a strong indicator of health. Generally, when coral tissue is healthy, the system can regulate potential overgrowth of bacteria and maintain a steady homeostasis. However, when a fragment is severely injured or dead, it is rapidly overrun and consumed by bacteria, a process fueled by a dramatic increase in oxygen consumption. In order to quantify this effect, we measured the 15-minute oxygen consumption profiles of healthy bleached corals (n = 59 fragments from N = 29 genotypes; positive control) and cryo-injured corals (n = 50, N = 24; injury induced by uncontrolled freezing in LN_2_; negative control), using a Loligo Systems temperature-controlled microplate respirometer (24 wells, 1700 μL volumes; Loligo Systems, Viborg, Denmark). The raw profiles for all samples were normalized to 100% O_2_ at time *t* = 0 and are shown in the first two panels of Fig. 3a. In order to facilitate comparison and statistical analysis, each profile was then fitted to an exponential function of the form *y* = *exp*(*-a*t*), characterized by the rate constant of oxygen consumption *a.* The distribution of this rate parameter describes the variability within each experimental group. The median values for the experimental groups (positive and negative controls) are depicted as the aggregate-fit lines in Fig. 3b, with the shaded regions denoting the 25% and 75% quartiles of *a.* These results demonstrate the characteristically different oxygen consumption profiles accompanying healthy and injured tissue, which we used as bounding markers of health in all additional experiments. Further details on fitting and all forthcoming statistical analysis can be seen in the Methods.

**Figure 3.**
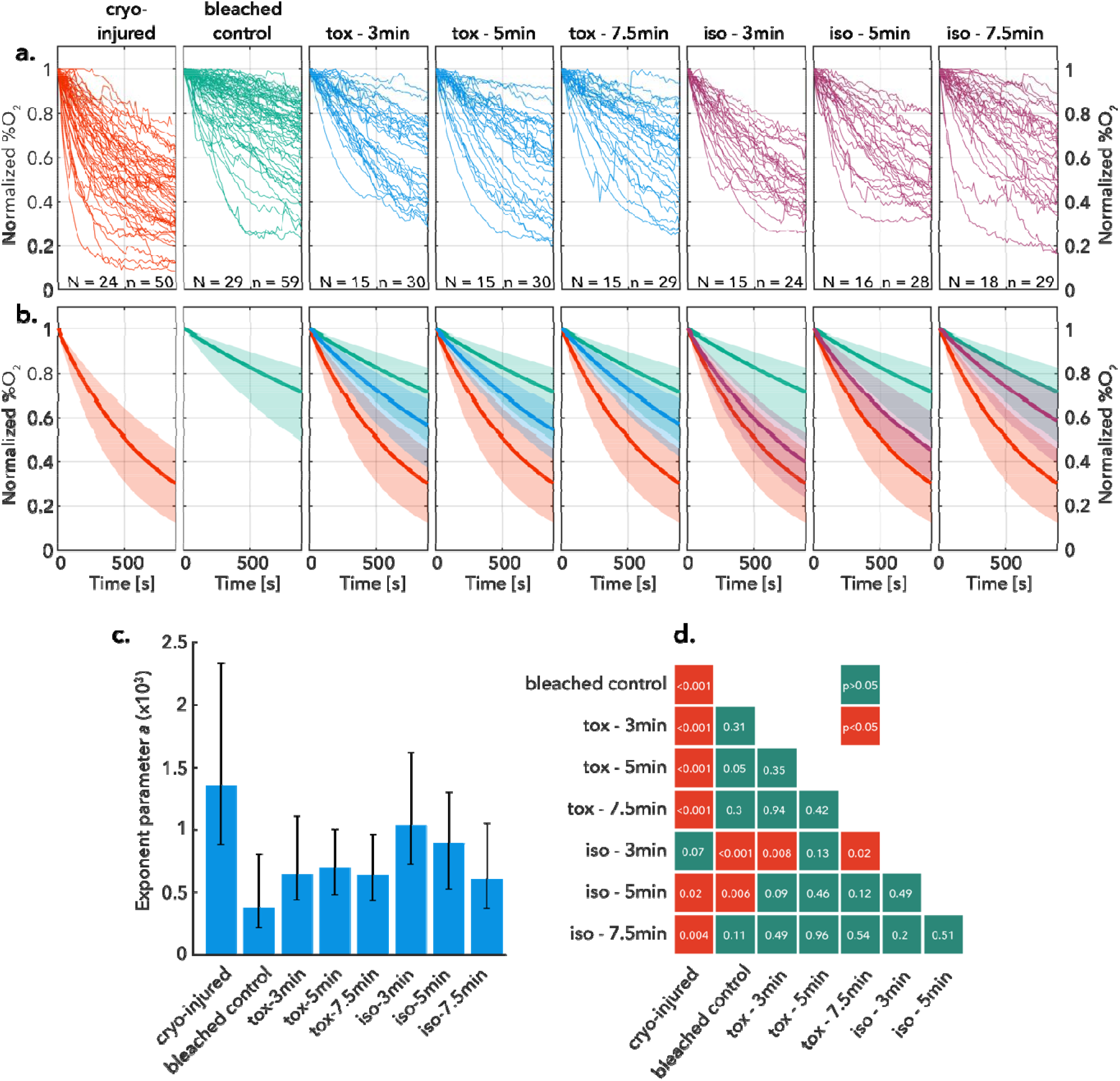
Coral respirometry results. (a) Raw and (b) exponentially-fitted normalized oxygen consumption curves for four groups of data: positive control (live bleached coral), negative control (cryo-injured coral), CVS1 toxicity experiments (labeled “tox – X”, where X is the coral fragment equilibration time in full-strength CVS1 cryoprotective solution), and isochoric vitrification experiments (labeled “iso -X”, where X is the same). Number of genotypes *N* and total samples *n* in each data set are displayed at the boom of each panel in (a). Solid lines in (b) represent median values and shaded regions are bounded by 25%/75% quartiles of the fitted exponential rate parameter *a*, which is itself plotted in panel (c) histogram with medians and 25%/75% quartiles of the exponential fit parameter for each treatment. Panel (d) provides a matrix of pairwise two-sample t-test results indicating significant difference between groups, demonstrating that isochoric vitrification after 7.5 minutes of osmotic equilibration in CVS1 produces coral that are statistically indistinguishable from healthy controls and statistically distinct from injured coral. *p*-values are displayed rounded to two decimal places.

### Experimental Pipeline

Having established the requisite husbandry to prepare mature coral for preservation and respirometry to evaluate their health in real-time thereafter, we designed an experimental pipeline to simultaneously probe for solution toxicity and cryo-injury in our model isochoric vitrification protocol. In order to ensure robust genotypic representation, for each genotype gathered, microfragments were dedicated to the following groups: positive control; negative control; toxicity trials; isochoric vitrification trials. In each of the toxicity and vitrification trials, three exposure times in full-strength CVS1 solution were tested (3, 5, and 7.5 min). The vitrification trials proceeded as described previously, and in the event that unintended nucleation was observed in the pressure signal (which was found typically to stem from issues related to the pressure transducer; discussed in later sections), the trial was marked and the fragment was segregated from those undergoing successful vitrification for the purposes of analysis. For the toxicity trials, the microfragments were transferred directly from the CPA loading stage shown in Fig. 2 to the rehydration stage, with the intention of isolating potentially deleterious chemical or dehydrative effects that may act independently of the cryopreservation process. Importantly, negative controls were cryo-injured by exposing them to precisely the same protocol as used for the vitrification trials, except without sealing the isochoric chamber, providing a sharp additional testament to the importance of isochoric conditions in facilitating the vitrification process.

### Coral fragments survive isochoric vitrification with sufficient osmotic conditioning

Fig. 3 showcases the aggregated respirometry profiles of these experiments, both in raw (a) and fitted (b) form, and the averaged oxygen consumption rate parameters ‘*a’* are shown in Fig. 3c. Pairwise two-sample t-tests were conducted for comparison between every group, and the resulting significance table is shown in Fig. 3d. Several important results emerge. First, all toxicity trials produced oxygen consumption profiles that are statistically indistinguishable from the positive bleached control, indicating that exposure to the CVS1 solution for any of the timepoints tested did not produce detectable damage on its own. Conversely, the vitrification trials showed a distinct dependence on CPA loading time; 3 min exposure (N=15, n=24) yielded respirometry profiles statistically similar to the negative controls (p = 0.07), indicating significant injury, and 5 min exposure (N=16, n=28) yielded profiles statistically different from both negative and positive controls (p = 0.02 and p = 0.006, respectively). Importantly, however, 7.5 min CPA exposure (N=18, n=29) yielded respirometry profiles both statistically indecipherable from the healthy controls (p = 0.11) and different from the negative controls (p = 0.004), indicating successful glass formation within the fragment and survival of the coral. To our knowledge, this represents the first successful cryopreservation and revival of the mature coral organism.

It is key to note further that all the data shown for the vitrification trials in Fig. 3 reflect successful vitrification of the external carrier solution surrounding the fragment, as indicated by the absence of a rise in pressure. Thus, the improvement in coral health post-vitrification with increasing CPA loading time suggests that, while the external solution may be vitrifying, ice formation may have been occurring *within* the coral tissue when the loading period was insufficient, i.e., at less than 7.5 min of equilibration time in full CVS1. This provides a testament to the essential role of effective dehydration and CPA loading for vitrification protocols, and as new techniques are developed to quantify CPA penetration in complex tissues, the degree of CPA penetration should be rigorously characterized. It should also be noted that, despite convective warming, no thermal stress cracking was observed in any samples.

### Developing an unmonitored, electronics-free cryobanking approach

Having identified a husbandry, loading, and vitrification pipeline that yielded surviving coral 24h post-thaw, we sought to modify our protocol to maximize its readiness for applied banking efforts. In our preliminary thermodynamic testing and applied preservation testing, continuous monitoring of the pressure was required to ensure no erroneous ice nucleation had occurred. However, continuous sample-to-sample monitoring with a pressure transducer is implausible for a real-world banking effort. The high-fidelity transducers required, not only add significantly to the cost of each preservation assembly, but also decrease the likelihood of successful vitrification, adding unnecessary additional surface area, material interfaces, opportunities for air entrainment, and thermal gradients to the system (all of which can help to stimulate ice nucleation)^37,39^. As such, in the final phase of our experiments, we eliminated the pressure transducer and all active monitoring / electronics from the system, in order to produce a more cryobank-ready preservation tool.

As shown in Fig. 4a, we replaced the pressure transducer and SS316 adapter with a solid Al7075 plug that employed the same metal-on-metal surface sealing mechanism and repeated our successful vitrification protocol (i.e., that with 7.5 min CPA equilibration time) on an additional set of coral microfragments (N=7, n =22). Leveraging the high statistical power of our respirometry results for injured coral, healthy coral, and coral deemed successfully vitrified by way of pressure verification, we then used the respirometry results of these 22 unmonitored trials to infer the success or failure of vitrification in the simple-capped system. As shown in Figs. 4b and 4c, the unmonitored trials yielded respirometry profiles statistically indecipherable from both the successful pressure-monitored trials and the control bleached trials (p = 0.15 and p = 0.63, respectively), suggesting both that this method yields continued successful vitrification and that this protocol may be used to the desired effect without any electronics or active monitoring.

**Figure 4.**
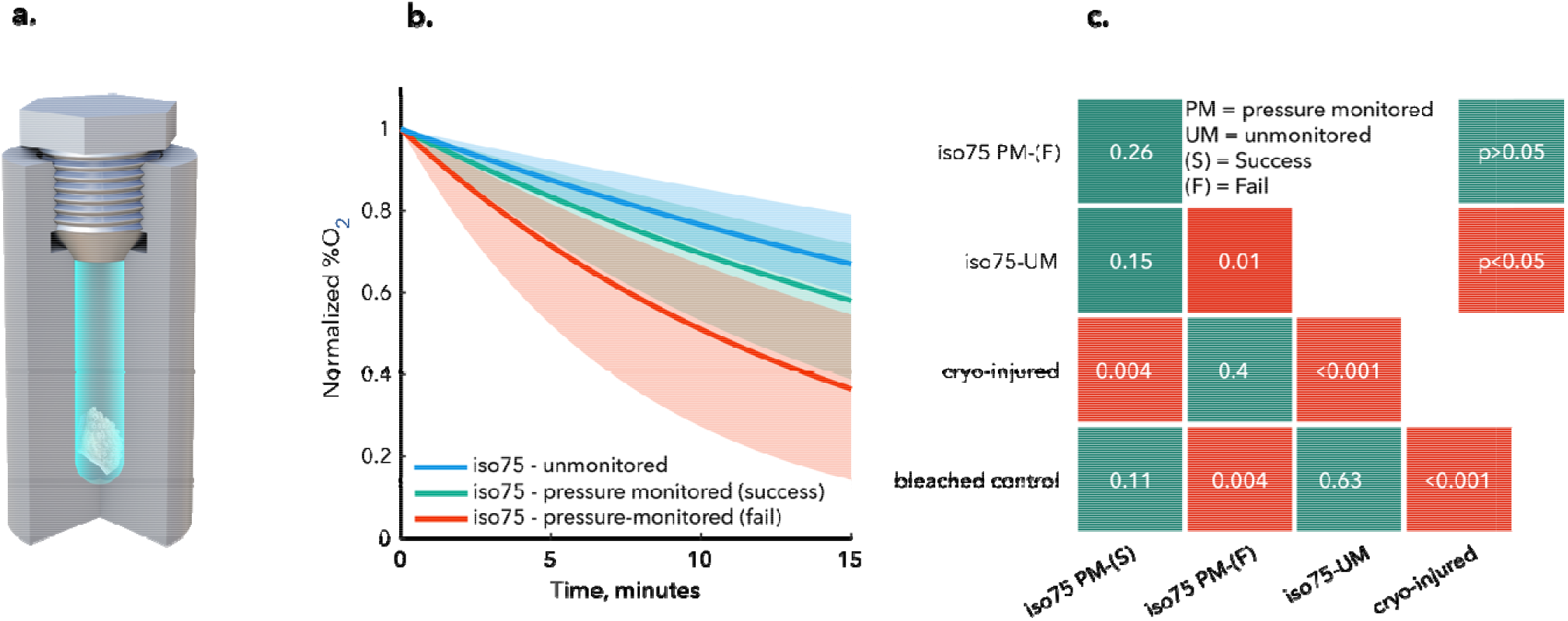
Unmonitored isochoric vitrification for scalable cryobanking. a) Passive, field-ready isochoric chamber that does require active monitoring or electronics of any kind. b) Aggregated median oxygen consumption curves for coral preserved via an unmonitored isochoric vitrification process vs. pressure-monitored equivalents deemed either successful or unsuccessful based on their pressure histories. Shaded regions are bounded by 25%/75% quartiles. c) Matrix of pairwise two-sample t-test results indicating significant difference between groups, showing that unmonitored samples are statistically indistinguishable from healthy bleached coral and statistically different from cryo-injured coral. Panels (b) and (c) also indicate that the unmonitored chamber shown in (a) produces better vitrification results in total as compared to the monitored device shown in Fig. 1a.

It should be further noted that removal of the pressure transduction apparatus appears likely to have increased the success rate of vitrification by potentially reducing the number of unintended nucleation events. We infer this by also comparing in Fig. 4b the respirometry results of the pressure-monitored trials that *failed* to vitrify (n = 9 out of a total of 37 trials). The unmonitored results in the capped system are statistically identical to those for healthy coral, whereas the pressure-monitored results that show nucleation are not. This suggests that the rate of successful vitrification is increased in the capped system relative to the measured 75.7% success rate in the aggregated pressure-monitored trials.

### Limitations of current data

The data presented in this work demonstrates a mL-scale isochoric vitrification process by which whole coral fragments may be cryopreserved, revived, and verified to be healthy at 24h post-thaw. However, significant further development of recovery methods and husbandry techniques will be required to move incrementally toward full restored health, resumed calcification, and long-term growth. Urgent topics of future research include potential effects of the cryopreservation process on the coral genome and microbiome; the efficacy of isochoric vitrification for other species of coral, which may not only have different biological responses but different propensities for stimulating ice nucleation based on their surface properties; and the ease of translation of isochoric vitrification into field conditions.

### Potential impacts of cryoconservation on the future of coral reefs

New models of future oceans that use an average temperature elevation of 1.5 °C^40–42^ suggest that only 2.5 to 5% of the worlds reefs may remain by the mid-2030s, restricted to two areas of the world. Given the limited scope of global efforts being made to mitigate heating of our oceans, the Intergovernmental Panel on Climate Change (IPCC) suggests that a future target of 2.0 °C and above is a more likely future scenario^42^, leaving a mere decade before the diversity of the world’s coral reefs may be all but lost.

Time is of the essence for coral reefs, and therefore we must take immediate, concrete steps to develop a wide suite of conservation practices with concomitant and robust *ex situ* processes. These processes include maintaining coral species in captivity^43^, creating coral mesocosm breeding facilities to facilitate and improve reliable breeding^44^, and enhancing existing biorepositories to include coral sperm, larvae, symbionts^11^, and coral fragments at critical global nodes^45^.

For *ex situ* cryostorage, sexually produced sperm and larvae are ideal given their size and assortment of genes, but this material is only available a few days per year and only in remote locations around the globe. Moreover, climate change and other stressors are increasingly and negatively impacting coral reproduction^13,46^ and, as a consequence, the physiological robustness of this produced sexual material.

If successfully transitioned into a restoration practice, isochoric vitrification could be applied strategically around the globe in relatively non-impacted areas hundreds of days per year, thereby quickly securing the genetic diversity and biodiversity of the whole coral organism using this field-friendly technology. Although coral microfragments are bleached before the isochoric process, they can be re-infected during the grow out process, and mature colonies can then be placed into *ex situ* reproduction facilities to yield sexually-produced offspring or transitioned to existing coral nurseries.

*Ex situ* processes will have a tremendous role to play in the future of maintaining coral reefs, and best practices mandate that these be applied side-by-side with current reef-based restoration practices that focus largely on *in situ* fragmentation and outplanting to current or former reefs. Isochoric vitrification provides a versatile example of one such process, and efforts to ready it for field applications should be undertaken with the utmost urgency.

## Methods

### Thermodynamic tests

In order to explore the thermodynamic premise of isochoric vitrification, chambers loaded with the CVS1 solution were cooled and warmed at different rates while the temperature and pressure of the system were monitored. *Chamber assembly and test preparation:* First, teflon tape was applied to both threaded sides of the 316 SS adaptor, which was then torqued into the ESI pressure transducer (rated for 0.1 to 400 MPa) to 27 ft-lbs. The CVS1 solution was loaded slowly into the 5.55 mL AL7075 chamber volume via syringe to avoid the formation of bubbles. The chamber was filled until the solution formed a convex meniscus over the top opening, then the adapter volume (0.22 mL) was filled with solution until a convex meniscus formed over the hole. The mated pressure sensor and adapter were torqued into the chamber to 45 ft-lbs, and excess solution was allowed to escape through the weephole. A type T thermocouple (OMEGA) was insulated with a layer of Kapton tape and inserted into the chamber weephole to monitor temperature throughout the experiment. The digital pressure transducer (Ellison Sensors Inc.) was wrapped in insulating bubble wrap to maintain the temperature above −50°C. A water warming bath fitted with two submersible water pumps (VIVOSUN) to provide turbulent mixing was warmed to 27°C. ESI-USB software and Omega Logging software were used to record the pressure and temperature, respectively. *Isochoric vitrification tests:* To achieve maximum cooling rates (96+/-6°C /min), the chamber was submerged up to the bottom of the pressure sensor throughout the process. Once nitrogen boiling ceased (indicating steady state), the chamber remained submerged for an additional two minutes before being rapidly moved to the warming bath. For maximum warming rates (387 +/- 55°C/min, n = 10), the chamber was submerged in the warming bath directly between the two pumps. The reduced cooling rates shown in Fig. 1c (4+/- 0.3°C /min, n = 10) were achieved by insulating the chamber with a plastic bag (P4M 4×6, GT Zip) before submerging in the liquid nitrogen. The reduced warming rates shown in Fig. 1d (4 +/- 0.1°C /min, n = 10) were achieved by removing the chamber from liquid nitrogen and allowing to cool in the air (at approximately 17 +/- 0.7°C). Between experiments, the chamber and adaptor volumes were rinsed with ethanol and vacuumed to remove any remaining liquid or particles. *Isobaric ice formation test:* To compare the effects of isobaric conditions on the CVS1 solution to those observed in the isochoric case, an unsealed chamber was used. The open AL7075 chamber was carefully filled with CVS1 solution up to the edge of the cylindrical chamber volume (not including the threaded region). The weephole of the chamber was sealed with Kapton tape to prevent the pooling of liquid nitrogen into the chamber. The chamber was then submerged up to the weephole in liquid nitrogen. The chamber was observed while it remained in the liquid nitrogen bath to avoid condensation on the surface. Ice growth was evaluated visually based on significant expansion of the solution.

### Coral colony selection and husbandry

From June – December 2022, small (approximately 10 cm x 20 cm) colony fragments of *Porites compressa* were selected from patch reef sites across Kāne□ohe Bay, O□ahu, Hawai□i and held in flow-through seawater mesocosms at the Hawai□i Institute of Marine Biology. Shortly after collection, colonies were processed into 0.5 cm^2^ microfragments (Page et al. 2018).

Microfragments were allowed to heal and recover for at least 10 days prior to use in experiments and were not used beyond 30days after microfragmentation. Corals were collected under Special Activity Permit Numbers SAP 2022-22 and SAP 2023-31 issued by the State of Hawai□i Department of Land and Natural Resources.

### Microfragment preparation

Due to differences in cryopreservation sensitivity between coral tissue and symbionts (Symbiodiniaceae require more time for cryoprotectant equilibration and faster freezing rates^34^), corals were bleached according to the protocol of Lager et al^33^. After 3-5 days of menthol and light exposure to induce bleaching, microfragments for each genotype were visually inspected under a dissecting microscope for tissue integrity and overall coloration. To ensure loss of symbionts, corals were then assessed for any photosynthetic activity (Yield) using an Imaging PAM (Heinz Walz GmbH, Germany). Microfragments that had visible tissue recession or photosynthetic yield values greater than 0.1 were not used in experiments. After visual and photosynthetic yield assessments, corals for each genotype were then assigned a treatment designation: bleached (live) control, dead (bleached negative control), toxicity controls (3 minutes, 5 minutes, and 7.5 minutes), and Isochoric (3 minutes, 5 minutes, and 7.5 minutes); the time designations corresponded to the length of time in the dehydrating cryoprotectant solution for both toxicity controls and cryopreserved specimens.

### Experimental treatments

Isochoric vitrification experiments and CVS1 toxicity experiments begin with the same procedure: portions of CVS1 were diluted with 0.22 micron filtered seawater (FSW) to 33% (10 mL FSW, 5 mL CVS1) and 66% (5 mL FSW, 10 mL CVS1) strength. Coral microfragments were then stepped into CVS1 at increasing concentrations: two minutes at 33%, two minutes at 66%, and 3, 5, or 7.5 minutes at 100% CVS1 depending on the experimental treatment designation. The isochoric chambers were filled with CVS1 per the same protocol used in the thermodynamic testing.

For toxicity trials, after completion of CPA equilibration, fragments were moved directly to the rehydration steps described below. For isochoric vitrification trials, fragments were placed using forceps into an isochoric chamber, taking care not to introduce any errant air bubbles.

Pressure-monitored vitrification trials then proceeded precisely as described in the thermodynamic testing section above. The unmonitored vitrification trials shown in Fig. 4 were performed on a subset of 7.5-minute corals, with the only difference in procedure being the replacement of the pressure transducer with an identically threaded solid cap.

After warming, the chamber was promptly opened and the microfragment was placed in a petri dish with 1 mL of 50% CVS1 for 30 seconds to begin rehydration. For the remainder of rehydration, FSW was added to the dish with increasing volumes over time: 1.8 mL after the initial 30 seconds in 50% CVS1, 3 mL after another 30 seconds, 9 mL after one minute, 18 mL after three minutes, and 20 mL after another three minutes. Toxicity controls and negative (cryo-injured) controls went through the same process of incremental dehydration in CVS1 and incremental rehydration.

Toxicity controls were not exposed to liquid nitrogen or warm bath. Negative controls were placed in an isochoric chamber and plunged in nitrogen according to the same procedure as described for the vitrification, but without sealing the chamber (with pressure sensor or solid cap), ensuring ice formation throughout the chamber.

### Respirometry Measurements

After isochoric vitrification, corals from all treatments were placed into new 6-well plates in 10 mL of 0.22 micron FSW and maintained at 26 °C with no light. After approximately 24 hours, a Loligo Microplate Respirometry system (Loligo Systems, Viborg, Denmark) was used to measure oxygen consumption as a proxy for coral health post-thaw. A small aquarium air pump was used to aerate a bottle of FSW to fill the 1.7 mL wells of the glass microplate with 26 °C oxygenated seawater for hydration of the microplate sensors, calibration of the unit (at high oxygen concentration), and experimental measurements. For low oxygen concentration calibration, 1 g of sodium sulfite was dissolved in 50 mL of deionized water and allowed to equilibrate for a few hours at 26 °C as oxygen-free water. For the respirometry measurements, aerated FSW and corals were placed in the wells of the microplate, and the microplate was sealed with PCR film, silicone pad, and compression block (provided in the Loligo Systems kit). Care was taken to ensure that no bubbles were sealed in the wells before starting the trial, and measurements were taken in 15 second intervals for 15 minutes. In addition to the sealed microplate and sensor, the respirometry system was assembled with a small aquarium pump and heater to circulate 26 °C FSW around the microplate in an acrylic chamber to maintain temperature for the duration of the assessments. The microplate wells were rinsed with deionized water between assessments.

### Respirometry Data Analysis

Respirometry measurements generated time series data of the oxygen concentration within each individual microplate well. These measurements were repeated on ≥20 samples for each experimental group. In order to draw conclusions from the variable responses, a data reduction pipeline was developed. First, the oxygen concentration measurements (%O2) were normalized to the initial concentration. Second, an exponential curve of the form *y=exp(-a*t)* was fit to the initial 5 minutes of normalized data using the *fit()* function in MATLAB 2022b, enabling each sample to be represented by a single exponential fit parameter *a*, and individual experimental groups to be represented by a distribution of the same. Statistical analysis of the respirometry data was conducted in order to compare the relative health of different experimental groups (e.g., cryo-injured, bleached control, toxicity, post-vitrification, etc.). The distributions of fitted exponential rate parameters computed for each group were compared using a two-sample t-test in MATLAB 2022b *(ttest2()* function). Effects are reported for significance levels of p < 0.05. A table of all exponential fit parameters is included in the SI.

## Data Availability

All data available upon reasonable request to the corresponding authors.

## Code Availability

All code (MATLAB) available upon reasonable request to the corresponding authors.

## Acknowledgements

This work was generously funded across institutions by a grant from the Revive & Restore Catalyst Science fund. The authors thank Andy Grams for graphic design help.

## Conflicts of Interest

BR filed a 2017 patent application related to isochoric vitrification, which is under review as of the date of submission of this work. MPP and BR have financial stake in a commercial entity that holds the license to said patent application. The other authors declare no conflicts of interest.

## Author Contributions

MPP and MH conceived the study as a whole and contributed all aspects. BR conceived the premise of isochoric vitrification. EMH, CL, RP, and KF carried out all experiments involving coral, with input from JD and MPP, and supervision from MH. BC performed all thermodynamic experiments, with support from ANC and supervision from MPP and BR. ANC performed the statistical analysis. MPP, EMH, MH, BC, and ANC wrote the paper, and all authors reviewed the manuscript and provided critical feedback.

## References

1. Bellwood, D. R., Hughes, T. P., Folke, C. & Nyström, M. Confronting the coral reef crisis. Nature 429, 827–833 (2004).

2. Bongaarts, J. IPBES, 2019. Summary for policymakers of the global assessment report on biodiversity and ecosystem services of the Intergovernmental Science-Policy Platform on Biodiversity and Ecosystem Services. Popul. Dev. Rev. 45, (2019).

3. Solomon, S. D. et al. Summary for Policymakers. In: Climate Change 2007: The Physical Science Basis. Contribution of Working Group I to the Fourth Assessment Report of the Intergovernmental Panel on Climate Changes. D Qin M Manning Z Chen M Marquis K Averyt M Tignor HL Mill. New York Cambridge Univ. Press pp Geneva, (2007).

4. Fisher, R. et al. Species Richness on Coral Reefs and the Pursuit of Convergent Global Estimates. Curr. Biol. 25, 500–505 (2015).

5. Bryant, D. L., Burke, D. L., McManus, J. & Spalding, M. Reefs at Risk: A Map-Based Indicator of Threats to the World’s Coral Reefs. (1998).

6. Elliff, C. I. & Silva, I. R. Coral reefs as the first line of defense: Shoreline protection in face of climate change. Mar. Environ. Res. 127, 148–154 (2017).

7. Spalding, M. et al. Mapping the global value and distribution of coral reef tourism. Mar. Policy 82, (2017).

8. Knowlton, N. et al. Rebuilding Coral Reefs: A Decadal Grand Challenge. (2021) doi: 10.53642/NRKY9386.

9. A decision framework for interventions to increase the persistence and resilience of coral reefs. A Decision Framework for Interventions to Increase the Persistence and Resilience of Coral Reefs (2019). doi: 10.17226/25424.

10. Hagedorn, M. et al. Assisted gene flow using cryopreserved sperm in critically endangered coral. Proc. Natl. Acad. Sci. U. S. A. 118, (2021).

11. Daly, J. et al. Cryopreservation can assist gene flow on the Great Barrier Reef. Coral Reefs 41, (2022).

12. Daly, J. et al. Successful cryopreservation of coral larvae using vitrification and laser warming. Sci. Rep. 8, (2018).

13. Hagedorn, M. et al. Potential bleaching effects on coral reproduction. in Reproduction, Fertility and Development vol. 28 (2016).

14. Baird, A. & Marshall, P. Mortality, growth and reproduction in scleractinian corals following bleaching on the Great Barrier Reef. Mar. Ecol. Prog. Ser. 237, 133–141 (2002).

15. Henley, E. M. et al. Reproductive plasticity of Hawaiian Montipora corals following thermal stress. Sci. Rep. 11, 12525 (2021).

16. Fahy, G. M., MacFarlane, D. R., Angell, C. A. & Meryman, H. T. Vitrification as an approach to cryopreservation. Cryobiology 21, 407–26 (1984).

17. Han, Z. & Bischof, J. C. Critical cooling and warming rates as a function of CPA concentration. CryoLetters vol. 41 (2020).

18. Zhan, L., Li, M. gang, Hays, T. & Bischof, J. Cryopreservation method for Drosophila melanogaster embryos. Nat. Commun. 12, (2021).

19. Khosla, K. et al. Successful Cryopreservation Of Zebrafish Embryos Followed By Hatching And Spawning With Laser Gold Nanowarming. Cryobiology 91, (2019).

20. Zhan, L. et al. Conduction Cooling and Plasmonic Heating Dramatically Increase Droplet Vitrification Volumes for Cell Cryopreservation. Adv. Sci. 8, (2021).

21. Zhan, L. et al. Pancreatic islet cryopreservation by vitrification achieves high viability, function, recovery and clinical scalability for transplantation. Nat. Med. 28, (2022).

22. Zhang, Y. et al. Isochoric vitrification: An experimental study to establish proof of concept. Cryobiology (2018) doi: 10.1016/j.cryobiol.2018.06.005.

23. Rubinsky, B., Perez, P. A. & Carlson, M. E. The thermodynamic principles of isochoric cryopreservation. Cryobiology (2005) doi: 10.1016/j.cryobiol.2004.12.002.

24. Powell-Palm, M. J., Rubinsky, B. & Sun, W. Freezing water at constant volume and under confinement. Commun. Phys. 3, (2020).

25. Powell-Palm, M. J., Koh-Bell, A. & Rubinsky, B. Isochoric conditions enhance stability of metastable supercooled water. Appl. Phys. Lett. 116, (2020).

26. Consiglio, A. N., Lilley, D., Prasher, R., Rubinsky, B. & Powell-Palm, M. J. Methods to stabilize aqueous supercooling identified by use of an isochoric nucleation detection (INDe) device. Cryobiology (2022) doi: 10.1016/j.cryobiol.2022.03.003.

27. Consiglio, A., Ukpai, G., Rubinsky, B. & Powell-Palm, M. J. Suppression of cavitation-induced nucleation in systems under isochoric confinement. Phys. Rev. Res. 2, (2020).

28. Sharma, A. et al. Cryopreservation of Whole Rat Livers by Vitrification and Nanowarming. Ann. Biomed. Eng. (2022) doi: 10.1007/s10439-022-03064-2.

29. Rubinsky, B. Mass transfer into biological matter using isochoric freezing. Cryobiology 100, 212–215 (2021).

30. Tyree, T. J., Dan, R. & Thorne, R. E. Density and electron density of aqueous cryoprotectant solutions at cryogenic temperatures for optimized cryoprotection and diffraction contrast. Acta Crystallogr. Sect. D Struct. Biol. 74, 471–479 (2018).

31. Rabin, Y. Mathematical modeling of surface deformation during vitrification. Cryobiology 102, 34–41 (2021).

32. Solanki, P. K. & Rabin, Y. Perspective: Temperature-Dependent Density And Thermal Expansion Of Cryoprotective Agents. Cryoletters 43, 1–9 (2022).

33. Lager, C. et al. Metrics of Coral Microfragment Viability. bioRxiv (2023) doi: https://doi.org/10.1101/2023.01.03.522625.

34. Hagedorn, M., Carter, V. L., Leong, J. C. & Kleinhans, F. W. Physiology and cryosensitivity of coral endosymbiotic algae (Symbiodinium). Cryobiology 60, 147–158 (2010).

35. Hagedorn, M. et al. Coral larvae conservation: Physiology and reproduction. Cryobiology 52, 33–47 (2006).

36. Perez, P. A., Preciado, J., Carlson, G., DeLonzor, R. & Rubinsky, B. The effect of undissolved air on isochoric freezing. Cryobiology 72, 225–231 (2016).

37. Huang, H., Yarmush, M. L. & Usta, O. B. Long-term deep-supercooling of large-volume water and red cell suspensions via surface sealing with immiscible liquids. Nat. Commun. (2018) doi: 10.1038/s41467-018-05636-0.

38. Camp, E. F. et al. The “Flexi-Chamber”: A Novel Cost-Effective In Situ Respirometry Chamber for Coral Physiological Measurements. PLoS One 10, e0138800 (2015).

39. Kar, A., Bhati, A., Lokanathan, M. & Bahadur, V. Faster Nucleation of Ice at the Three-Phase Contact Line: Influence of Interfacial Chemistry. Langmuir 37, 12673–12680 (2021).

40. Kalmus, P., Ekanayaka, A., Kang, E., Baird, M. & Gierach, M. Past the Precipice? Projected Coral Habitability Under Global Heating. Earth’s Futur. 10, (2022).

41. Dixon, A. M., Forster, P. M., Heron, S. F., Stoner, A. M. K. & Beger, M. Future loss of local-scale thermal refugia in coral reef ecosystems. PLOS Clim. 1, e0000004 (2022).

42. Intergovernmental Panel on Climate Change (IPCC). The Ocean and Cryosphere in a Changing Climate. (Cambridge University Press, 2022). doi: 10.1017/9781009157964.

43. Zoccola, D. et al. The World Coral Conservatory (WCC): A Noah’s ark for corals to support survival of reef ecosystems. PLOS Biol. 18, e3000823 (2020).

44. Craggs, J. et al. Inducing broadcast coral spawning ex situ: Closed system mesocosm design and husbandry protocol. Ecol. Evol. 7, 11066–11078 (2017).

45. Bouwmeester, J., Daly, J., Zuchowicz, N. & Hagedorn, M. Cryopreservation to Conserve Genetic Diversity of Reef-Building Corals. in 225–240 (2022). doi: 10.1007/978-3-031-07055-6_14.

46. Levitan, D., Boudreau, W., Jara, J. & Knowlton, N. Long-term reduced spawning in Orbicella coral species due to temperature stress. Mar. Ecol. Prog. Ser. 515, 1–10 (2014).

