## Supplementary Information for "Cryopreservation and revival of Hawaiian stony corals via isochoric vitrification"

*^4^Hawaiʻi Institute of Marine Biology, University of Hawaiʻi at Mānoa, Kāneʻohe, HI, 96744, USA*

### S0. CVS1 Solution Design

CVS1 was based on a previous solution developed for the vitrification and laser-assisted re-warming of droplets containing coral larvae^1^. The original formulation contained DMSO (1M) Propanediol (1M) Glycerol (1M) and Trehalose (1.3M). This concentration of trehalose is supersaturated at low-temperature however, and while previous droplet experiments remained stable through thermal cycling, at mL-scale volumes solid trehalose was found to precipitate out of solution. Thus, for CVS1, the concentration of trehalose was reduced to ~0.85M, which was found to successfully avoid unintended precipitation. The concentration of each of the small molecules was also increased by approximately 0.05M in order to compensate for the reduced trehalose concentration.

We should note that, while molarity is one of the most commonly reported concentration metrics in the vitrification literature, it is not as rigorous as molality or weight percent, which do not depend on environmental conditions (e.g., standard temperature and pressure). Thus, we also provide the CVS1 recipe in mass percent below.

***Table S1.*** *CVS1 Recipe*

|  | $x$ (w/w) | Approx. molarity (M) |
| --- | --- | --- |
| PBS | 0.504 | 0.15 |
| DMSO | 0.074 | 1.05 |
| PG | 0.072 | 1.05 |
| Glycerol | 0.087 | 1.05 |
| Trehalose | 0.263 | 0.85 |

### S1. COMSOL Cooling / Warming Rate Estimations

Due to physical limitations of the sealed isochoric chamber, the thermocouple temperature measurements are made near the inner edge of the chamber wall. Thus, the cooling and warming rates measured by the thermocouple represent the maximum possible values experienced in the solution, i.e., at the interface of the aluminum and the solution. A simple cylindrical 1D axisymmetric thermal transport model was created in COMSOL Multiphysics in order to estimate the cooling and warming rates at the *center* of the solution volume, which represent the *minimum* rates, under the experimentally observed peripheral temperature conditions. Properties of pure water were used, and a simple cooling and warming path was constructed using average values from the vitrification experiments.

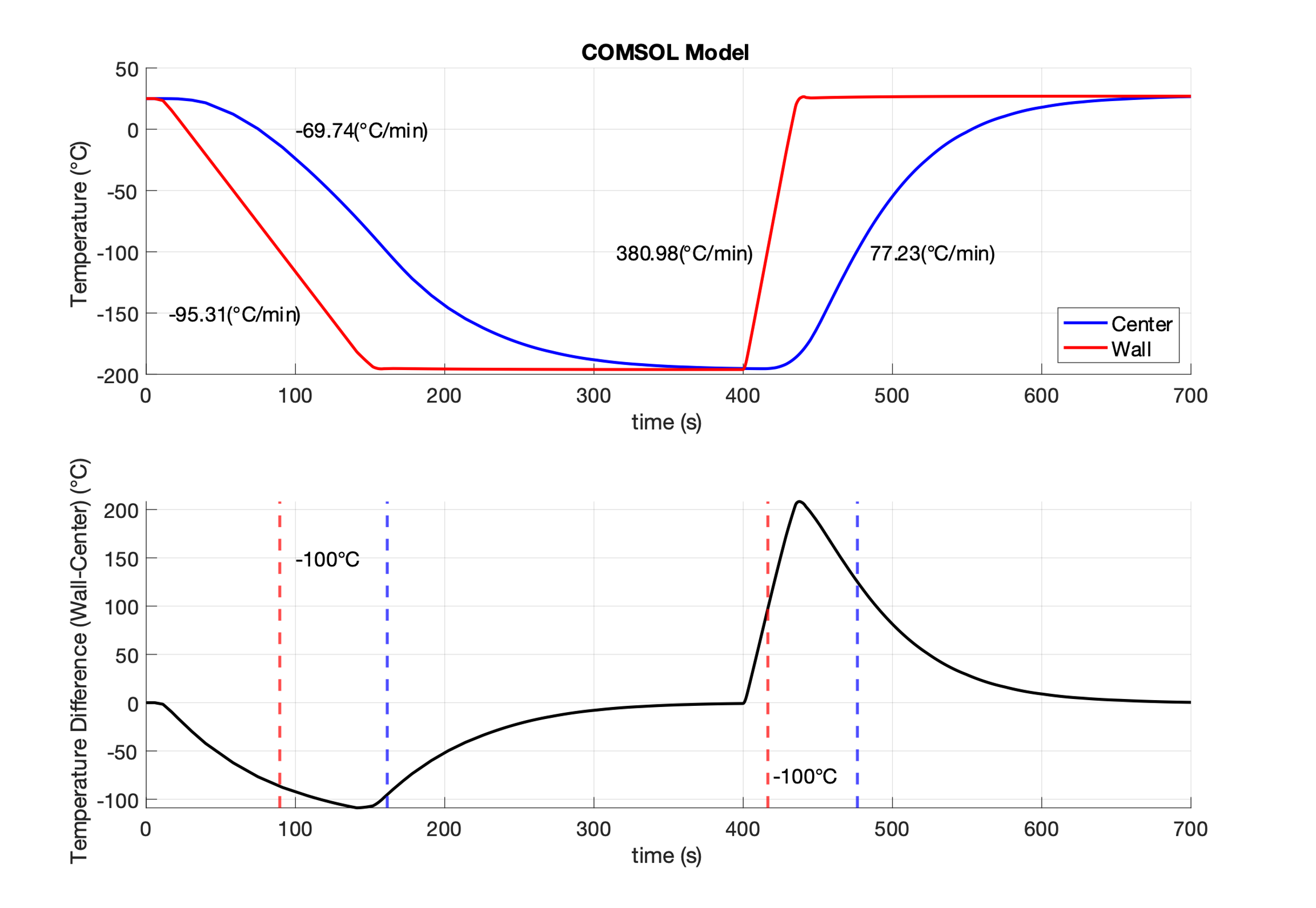

**Figure S1**. COMSOL results. The top panel shows the COMSOL model of the estimated temperature at the wall (red) and at the center of the solution (blue), with average cooling and warming rates noted. The bottom panel shows the temperature difference between the two locations (Wall Temperature – Center Temperature) with vertical dashed lines indicating the time point at which the temperature curves reach -100°C.

The results of the simulation are seen in Figure S1. Average cooling and warming rates were estimated for temperatures between -100°C and 0°C, representing the temperature range above the glass transition in which ice nucleation is possible. While the cooling and warming rates of the fluid at the inner wall of the chamber are nearly the same as for the chamber (~95°C/min and ~380°C/min, respectively), the average cooling rate at the center axis is ~70°C/min and the average warming rate at the center axis is ~77°C/min. These results highlight both the difficulty of achieving fast cooling/warming rates in bulk systems and the exceptional capacity of CVS1 to vitrify at slow rates under isochoric conditions.

When convectively cooling and warming a system during vitrification and rewarming, large thermal gradients can form. These thermal gradients may generate stresses in the vitrifying sample due to non-uniform thermal expansion and have been observed to result in mechanical fracture of the vitrified material^2,3^. It is difficult to quantify the precise conditions that result in thermal stress cracking due mainly to uncertainties in the thermal and mechanical properties of the vitrifying material, however experiments performed in open systems at bulk volumes with critical dimensions of ~1-2cm have observed cracking when convectively rewarmed^4,5^.

In order to quantify the potential for thermal stress cracking, vitrification studies often report the temperature difference between the outside and center of the vitrifying sample. Depending on the material, critical temperature differences are often reported as ~20-50°C^4,5^. Figure S2 shows the temperature difference between the fluid at the center axis of the chamber and fluid near the inner wall of the chamber computed from the results of the heat transfer simulation. While this simulation is highly simplified (only axial conduction and using water thermal properties), we find temperature differences >100°C. We should note that thermal stresses are in reality a function of thermal gradient (i.e., dT/dx) and not of the temperature difference between two arbitrary locations. Thus, while we can compute the center-wall temperature difference from the simulation, it is not completely possible to compare this to data from other vitrification experiments.

The large thermal gradients computed from the simplified thermal model suggest that there is a high potential for thermal stress cracking in the isochoric vitrification experiments. Interestingly however, we observe no evidence of stress cracking during the isochoric vitrification experiments. Whether this is a result of the isochoric confinement or the coral specimen’s mechanical toughness should be investigated in future studies. We will note however that studies have shown that stress cracking during vitrification typically stems from massive deformation of the open liquid-air interface, which contracts dramatically. No such open boundary exists in isochoric systems, and this may contribute to enhanced cracking resistance in isochoric systems.

### S2. Glass transition temperature

The model of Couchman and Karasz^6,7^ provides a method to approximate the glass transition temperature of aqueous solutions based on the relative component mass fraction and the glass transition temperature of the pure components.

$$\frac{1}{T_{g,solution}}=\sum_{i} \frac{x_{i}}{T_{g,i}}$$

Here, $T_{g,solution}$ is the glass transition temperature of the solution [K], $x_{i}$ is the $i$-th component mass fraction in solution, and $T_{g,i}$ is the glass transition temperature of the $i$-th component [K]. The table below shows the glass transition temperature and solute mass fractions for the CVS1 solution. From this data, we can calculate an approximate glass transition temperature of –100°C.

***Table S2.*** *Glass transition temperatures of CVS1 components*

|  | $x$ (w/w) | $T_{g}$ (K) | reference |
| --- | --- | --- | --- |
| PBS | 0.504 | 136 | value for water used^8^ |
| DMSO | 0.074 | 141 | ^9^ |
| PG | 0.072 | 167 | ^10^ |
| Glycerol | 0.087 | 190 | ^10^ |
| Trehalose | 0.263 | 373 | ^11^ |

### S3. Partial vitrification

When cooled at a rate slower than the critical cooling rate for vitrification, ice nucleates and grows within the system. This process pressurizes the isochoric system and leads to ripening of the liquid (i.e., increase of its solute concentration). By estimating that 50% of the water in the system crystallizes into ice, we can approximate the amount by which the concentration of the liquid has increased. These concentrations are shown in Table S3.

The glass transition temperature for a solution of this composition can be estimated by again employing the Couchman and Karasz relation. We obtain a value of –85°C. This temperature closely coincides with the plateauing of the pressure trace during the slow cooling trial in Figure 1c of the main text. After the freeze concentrated solution vitrifies, all ice growth ceases, and the system begins to strongly contract as the temperature is progressively lowered.

***Table S3.*** *Freeze concentration of CVS1*

*components if 50% of water were to freeze.*

|  | $x$ (w/w) |
| --- | --- |
| PBS | 0.335 |
| DMSO | 0.099 |
| PG | 0.097 |
| Glycerol | 0.117 |
| Trehalose | 0.352 |

### S4. Intermediate Cooling Rate Experiments

The thermodynamic tests reported in the main text were conducted with fast cooling rates of ~96°C/min and slow cooling rates of ~4°C/min. These experiments were repeated for intermediate cooling rates of 12°C /min, 43°C /min, and 85°C /min (various insulating materials were wrapped around the chamber to achieve the desired cooling rates). The pressure and temperature traces for these experiments are shown in Figure S2.

We observe that the maximum pressure decreases as the cooling rate increases. This trend may be evidence that at higher cooling rates, a smaller fraction of the system freezes which results in a larger portion of the solution vitrifying.

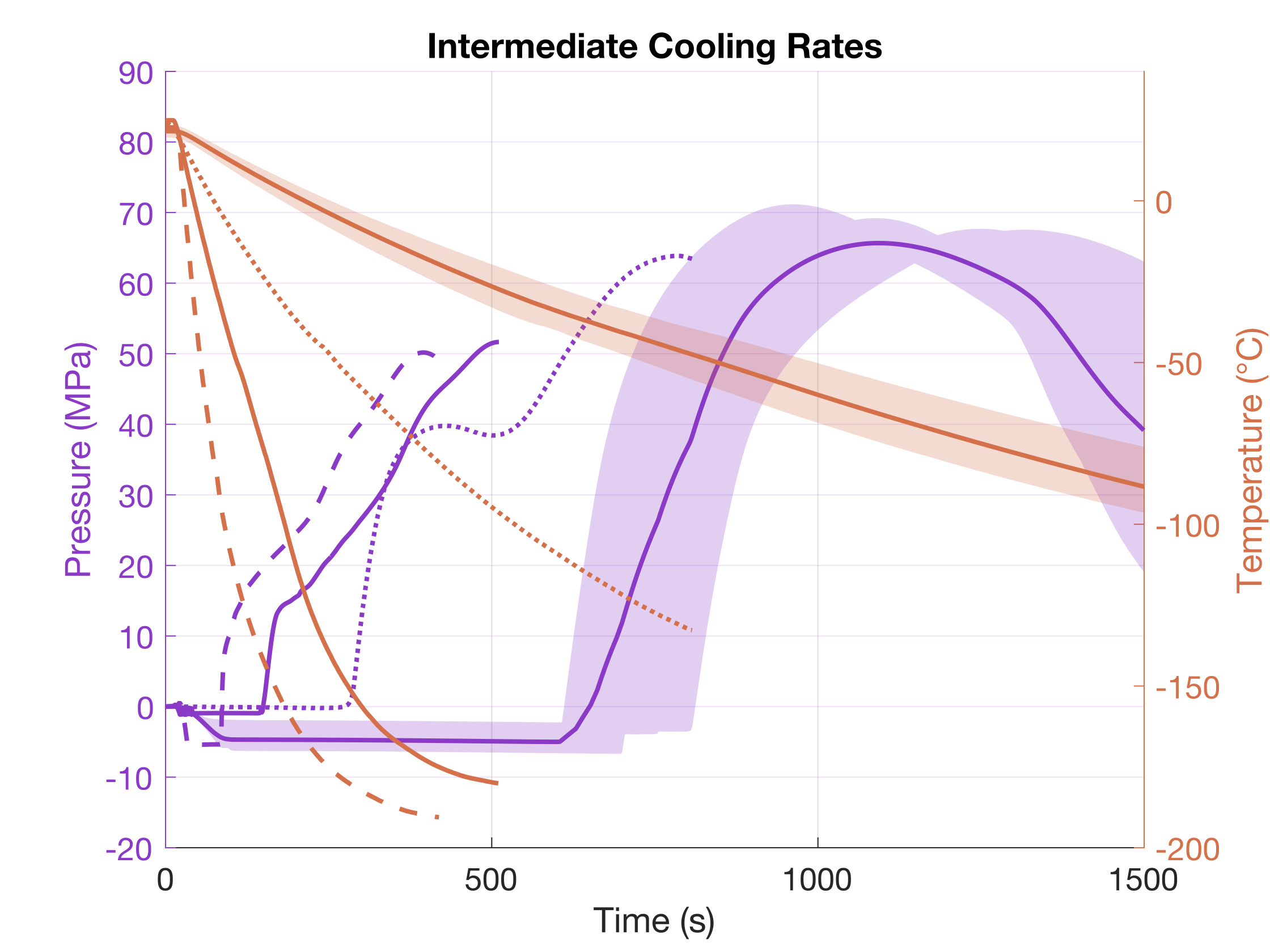

**Figure S2**. PTT traces for slow (4+/- 0.3°C /min, n = 9, shown in the solid, shaded lines) and intermediate cooling rates (12°C /min in the dotted lines, 43°C /min in the solid, unshaded lines, and 85°C /min in the dashed lines). The decreasing peak pressure accompanying increasing cooling rates suggests that a greater degree of partial vitrification may be occurring as cooling rates increase, until the threshold at which complete vitrification occurs (estimated to be ~96 °C/min).

### S5. Isobaric vitrification experiments

In order to evaluate the effect of isochoric confinement on the vitrification process, we repeated the vitrification experiments without the top plug. Figure S3 shows images of the unsealed chamber filled with CVS1 and plunged into nitrogen. Ice formation was observed in each of the experiments as evidenced by the visible protrusion of the crystallized solution above the original fill line. This expansion is indicative of ice formation since complete vitrification typically results in net contraction of a solution^12–14^.

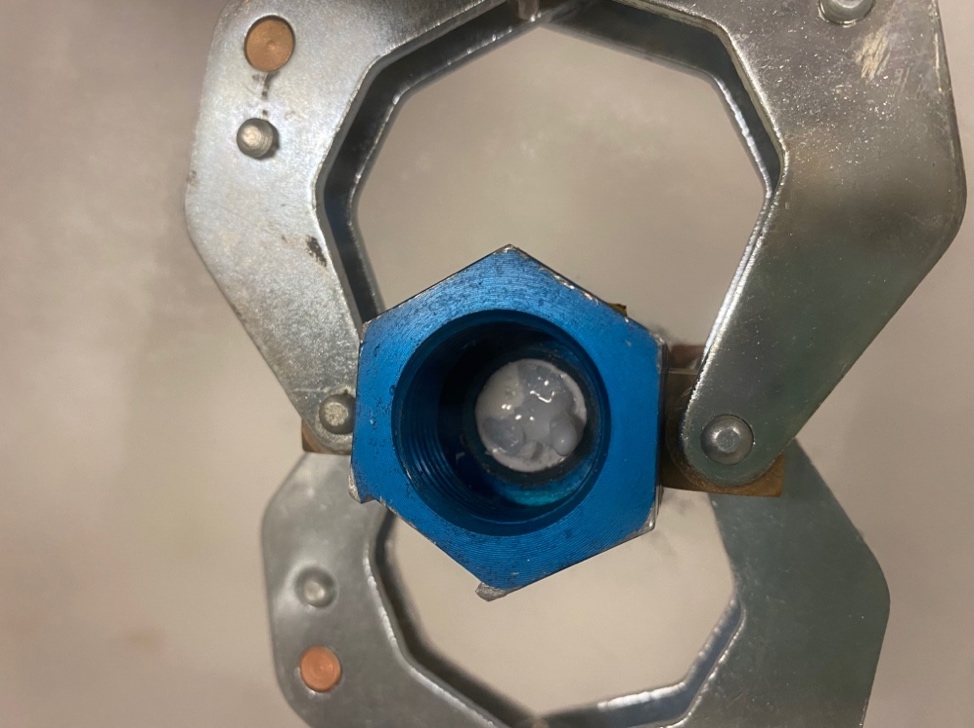

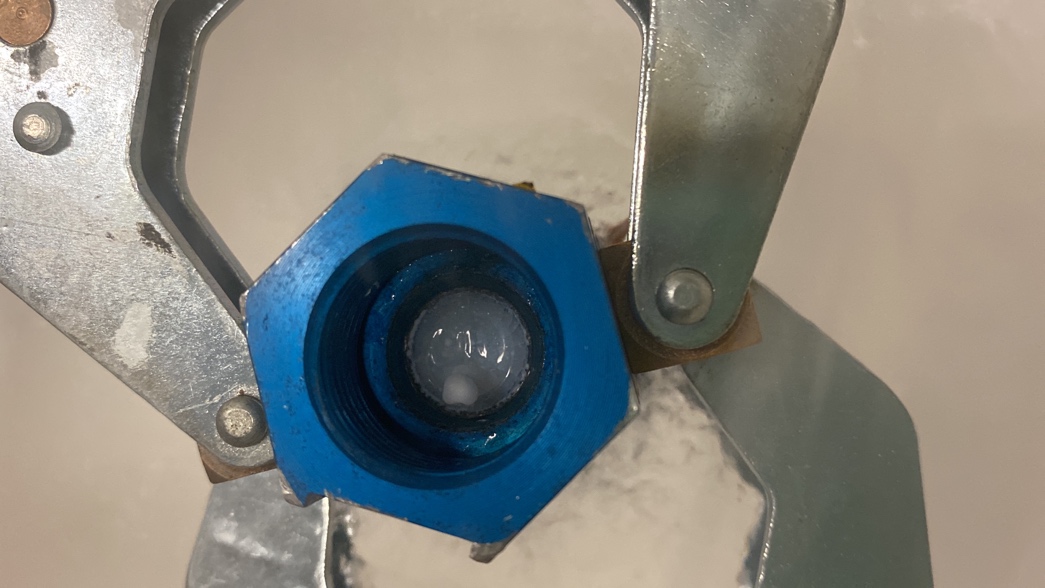

**Figure S3.** Images of two isobaric experiments performed with the CVS1 solution in an unsealed chamber. Images were taken while the chamber remained in the liquid nitrogen to avoid condensation. Ice nucleation is evidenced by the protruding solidified mass.

### S6. Isochoric Vitrification of *Porites compressa* Fragments – Detailed Procedure

- Begin with healthy (good tissue integrity) bleached coral microfragments (approx. 1.5 cm^2^) that have rested at least 10 days to heal since fragmentation process
  - For information on coral microfragmentation, see Page et al. 2018^15^
  - For information on coral bleaching with menthol and associated health metrics, see Lager et al. 2023^16^

*Setup and preparation*

- Prepare 200-300 mL of CVS1 cryoprotectant solution and 2 liters of 0.22-µm filtered seawater (FSW)
  - If CVS1 has been stored in the fridge overnight, warm to room temp and mix thoroughly before use – no precipitate should be present and avoid microbubble formation
- Wrap threads of isochoric chamber adapter (if using pressure sensors) or solid chamber cap (if not using pressure sensors) with Teflon tape
  - If using pressure sensors, thread adapter onto sensor and tighten to 27 ft.-lbs.; wrap and secure pressure sensor with insulation (bubble wrap, flexible foam, or similar).
- Place chamber in vice, and using a syringe with a 6 cm needle, slowly fill the chamber with 100% CVS1
  - Start at bottom and slowly pull syringe up while filling to ensure that no air bubbles form
    - Chamber should be filled to the point where the threads meet the interior cavity
    - This typically requires ~8 mL of CVS1
  - Recheck interior cavity of chamber for any bubbles and remove if present with syrunge
- Dehydration Station (set-up ahead of time before vitrification)
  - In two 15 mL tubes, prepare dilutions of 33% (10 mL FSW, 5 mL CVS1) and 66% (5 mL FSW, 10 mL CVS1)
  - Label and fill three small dishes each with 10 mL of 33%, 66%, and 100% CVS1
- Rehydration Station (set-up ahead of time for after vitrification)
  - Set up 27 °C warm bath with circulating water
    - Submersible pumps can be used to vigorously circulate water
  - Fill a 15 mL tube with 50% cryoprotectant dilution (7.5 mL FSW, 7.5 mL CVS1)
    - Fill a large Petri dish with 1 mL of 50% CVS1
  - Fill four 15 mL tubes with 3 mL, 9 mL, 18 mL, and 20 mL of FSW
  - Prepare one 50 mL tube with FSW
- Fill a cryo-safe tub with enough liquid nitrogen to submerge an isochoric chamber (not including the attached sensor) without the chamber contacting the bottom
  - Cover until ready to use to reduce loss of liquid nitrogen

*Stepping corals into CVS1 solution*

- Using forceps, remove microfragment from seawater and dab on Kimwipe 2-3 times to remove as much seawater from fragment as possible (do not damage the tissue)
- Place fragment in 33% CVS1 for 2 minutes
  - Gently pipette solution around coral a few times
  - Remove fragment and dab on Kimwipe
- Place microfragment in 66% CVS1 for 2 minutes
  - Gently pipette solution around coral
  - Remove fragment and dab on Kimwipe
- Place fragment in 100% CVS1 dish for 1 minute, pipette solution around fragment, and place in hex chamber filled with 100% CVS1 solution
  - The two minutes in 33% and 66% CVS1 *do not count* toward the target dehydration time (3 min, 5 min, or 7.5 min) – only when corals are placed into 100% CVS1
- The solutions in the three dishes of 33%, 66%, and 100% can be reused but should be replaced after 3-4 corals have passed through

*Isochoric vitrification*

- Using a syringe, fill the adapter attached to the sensor with 100% CVS1. Insert the tip to the very bottom of the cavity and fill slowly, gradually pulling the syringe while dispensing solution.
  - *If using the solid chamber cap and no pressure sensor with adapter, skip this step*
- With the hexagonal isochoric chamber secured in a vice, wrap a paper towel around the chamber weep hole and thread adapter with pressure sensor (or cap if no sensor is present) to the hex chamber and hand tighten – then tighten to 45 ft-lbs. with torque wrench
- Attach pressure sensor to USB cable and begin recording the pressure signal once the fragment has been exposed to 100% CVS1 for the allotted treatment time point
  - *Skip this step if no pressure sensors are used*
- Submerge isochoric chamber into liquid nitrogen
  - Hold the chamber such that the top of the chamber is below the surface of the liquid nitrogen and *does not contact the bottom* of the container, leaving the sensor in air just at the surface of the liquid nitrogen (LN_2_)
    - If using a solid chamber cap (no sensor), secure the chamber to a sturdy metal wire or similar with enough length to handle above LN_2_ and submerge until cap is covered by LN_2_
  - Hold sensor and chamber steady while temperature equilibrates (until LN_2_ boil)
    - After the LN_2_reaches a final boil, leave the chamber suspended in LN_2_ for two additional minutes
  - To thaw the unit, quickly transfer sensor and chamber to the 27°C warming bath for 2 minutes
    - As in cooling, submerge the chamber into the water, leaving the sensor exposed to air, without allowing the chamber to contact the bottom of warm bath
      - If no sensor, submerge chamber and cap below water line
    - The water should be vigorously circulating around all sides of the chamber
- Once thawed, secure chamber in vice and remove pressure sensor/adapter assembly (or cap) and pour solution with fragment into holding dish

*Rehydration – Stepping corals out of CVS1 solution*

- Place fragment in Petri dish with 50% CVS1 for 30 seconds, gently pipette solution around coral, then add 1.8 ml of FSW and gently pipette to mix
  - After another 30 sec (!-min post-thaw), add 3 mL FSW and gently pipette to mix gently
  - After 1-min (2-min post-thaw), add 9 mL FSW and mix gently
  - For both 3-min and 5-min dehydration treatments, add 18 mL and 20 mL FSW in 1-min increments (3- and 4-minutes post-thaw, respectively), gently mixing at each step
  - For 7.5-min dehydration, add 18 mL and 20 mL FSW in 3-min increments (5- and 8-min post-thaw, respectively), gently mixing at each step
  - Regardless of dehydration treatment time, allow coral to sit in petri dish for another minute
- Place coral back into 10 mL FSW in 6-well plate (or in desired holding tank)

*Toxicity and Cryo-damaged (negative) controls*

- For toxicity controls (Tox 3-min, Tox 5-min, and Tox 7.5-min) follow the same step in dehydration and rehydration process, as described above, but skip the liquid nitrogen (freezing) and warm bath steps, moving from 100% CVS1 (after 3-, 5-, or 7.5-min) to the 50% CVS1 rehydration phase, as described above
- For Cryo-damaged (negative) controls, gradually step corals into CVS1 as described above (typically the 5-min or 7.5-min treatment) and into an open hex-chamber of 100% CVS1
  - *Do not cap the chamber*
  - Instead, submerge the chamber in LN_2_ until it is a few mm above the liquid
  - Intermittently move forceps from LN_2_ into contact with the CVS1 solution in the chamber, stimulating early nucleation (ice formation) to occur
  - Allow for LN_2_ boil, wait two minutes, then warm and rehydrate per above methods, keeping the lip of the chamber above the water level, and place into 10 mL FSW in well plates

*Respirometry (using Loligo Systems Microrespirometry)*

- Prepare at least 2.5 liters of 0.22-µm FSW
- Fill one plastic squirt bottle with FSW and another with deionized (DI) water
  - Using a small air pump with air stone, bubble air into the FSW bottle
- Prepare a 50 mL tube of DI with 1.2 – 1.3 g of sodium sulfite for oxygen-free water
  - Dissolve Na_2_SO_3_ in DI, equilibrate to 26°C, and let stand for 2-3 h before using
  - *This is separate from the regular (“bulk”) DI bottle above*
- Set up all sensors and accessories required to read respirometer (per Loligo Systems instructions)
- To maintain the temperature constant in the Loligo chambers throughout the experiment, fill small aquarium tank with enough FSW to cover a small aquarium heater and powerhead and to also allow water to cycle through acrylic respirometry chamber (approx. 1.5 – 2 liters of FSW)
  - Set temperature to 25-26°C and monitor with thermometer
- Fill wells of the glass respirometry sensor Loligo plate with bubbled FSW, allowing the oxygen sensors on the bottom of the glass plate to hydrate in oxygenated FSW for at least 30-45 minutes at the proper temperature (25-26 °C)
- Calibration of sensor (taken from manufacturer’s instructions)
  - Ensure that all settings on respirometry software correspond with experiment (e.g., salinity, temperature, etc.)
  - Fill wells with bubbled filtered seawater, allow software to read the wells, and input calibration as “High” range (wells should read at or near 100% air saturation)
    - Drain and rinse with DI water
  - Fill wells with oxygen free (DI-sodium sulfite) water and ensure that wells are reading near 0% air saturation (anything less that 10% is fine)
    - In settings, change salinity from 35 PPT to 0 PPT
    - Input calibration as “Low” range
    - Change salinity back to 35 PPT
  - Save file
- Assign each well to be used a coral treatment name, “Blank” (empty well for background microbial activity), or “Not Set” (not used)
  - Skeleton (skeleton of coral fragment that has been bleached, rinsed, and hydrated in FSW) – can be used in place of ‘Blank’
  - Bleached coral fragment – Live control
  - Dead control – Fragments frozen in under icobaric conditions with ice formation
  - Tox 3, 5, and 7.5 min– Controls for freezing process
  - Iso 3, 5, and 7.5 min – Experimental treatment times
  - IsoXCap
    - Where ‘X’ is dehydration time and no pressure sensor was used (e.g., Iso7.5Cap)
- Rinse wells with DI and fill with bubbled FSW, allow software to read bubbled FSW two or three times, and then normalize wells to 100% (feature in respirometry software)
- Load fragments into respective wells
  - Seal plate with silicone pad and block (included in kit) and ensure that no air bubbles have formed in the wells of the plate
  - Place sealed plate into water bath and confirm that it is seated properly
  - Start time log and timer
  - Seal top of water bath
  - Place unit on shaker plate to gently move unit while running
    - If not in possession of a shaker plate, manually agitate unit approximately every minute to mix water in wells
  - Sensors will take a reading every 15 seconds with default software settings
    - Run log for 15 minutes
- Take corals out of wells and place back into their respective 6-well plates
- Rinse glass respirometry plate with DI water before next use
- Once finished breakdown equipment and rinse in fresh water
- Fill wells of glass sensor plate with 50/50 bleach solution (half household Clorox^TM^ bleach, half DI water), let stand for 10 minutes, rinse several times with DI, and store in black manufacturer bag

### S7. Coral fragment oxygen consumption exponential rate parameters *a* for all treatments

| Sample  Number | skeleton | cryo-injured | bleached control | tox3 | tox5 | tox7.5 | iso3 | iso5 | iso7.5  monitored success | iso7.5  unmonitored | iso7.5  monitored  fail |
| --- | --- | --- | --- | --- | --- | --- | --- | --- | --- | --- | --- |
| 1 | 3.91E-07 | -2.83E-05 | -1.40E-05 | -2.03E-06 | -4.94E-06 | -2.92E-06 | -8.26E-06 | -1.05E-05 | -3.05E-05 | -4.53E-06 | -1.88E-05 |
| 2 | 1.46E-07 | -1.23E-05 | -3.94E-06 | -1.19E-05 | -1.76E-06 | -8.25E-06 | -7.11E-06 | -9.96E-06 | -5.79E-06 | -1.35E-05 | -4.34E-06 |
| 3 | 4.98E-07 | -7.73E-06 | -1.97E-06 | -2.23E-06 | -3.97E-06 | -5.41E-06 | -1.38E-05 | -7.29E-06 | -6.82E-06 | -3.33E-06 | -2.70E-05 |
| 4 | -1.56E-07 | -1.07E-05 | -6.39E-06 | -6.04E-06 | -4.52E-06 | -6.01E-06 | -8.06E-06 | -3.94E-06 | -7.98E-06 | -1.66E-06 | -9.92E-06 |
| 5 | 7.10E-07 | -1.65E-05 | -8.67E-06 | -5.17E-06 | -1.68E-05 | -1.95E-06 | -2.23E-05 | -5.14E-06 | -5.43E-06 | -2.08E-06 | -6.81E-06 |
| 6 | 1.09E-06 | -3.74E-05 | -2.05E-05 | -6.96E-06 | -7.36E-06 | -2.57E-06 | -1.12E-05 | -6.00E-06 | -2.54E-05 | -2.43E-06 | -7.10E-06 |
| 7 | 2.55E-07 | -2.48E-05 | -1.78E-07 | -4.77E-06 | -7.18E-06 | -6.64E-06 | -5.48E-06 | -1.86E-05 | -1.00E-05 | -5.74E-06 | -1.17E-05 |
| 8 | 7.26E-07 | -1.07E-05 | -2.82E-06 | -4.87E-06 | -5.69E-06 | -6.85E-06 | -6.21E-06 | -3.27E-06 | -2.06E-06 | -1.69E-06 | -1.67E-05 |
| 9 | 1.04E-06 | -5.78E-06 | -2.54E-06 | -4.89E-06 | -4.89E-06 | -4.24E-06 | -9.37E-06 | -4.35E-06 | -3.66E-06 | -4.22E-06 | -2.87E-06 |
| 10 | 7.35E-07 | -2.70E-05 | -1.95E-06 | -1.28E-06 | -3.54E-06 | -1.78E-06 | -9.10E-06 | -1.83E-05 | -1.13E-05 | -5.42E-06 |  |
| 11 | 1.93E-06 | -2.74E-05 | -2.48E-06 | -1.59E-06 | -1.84E-06 | -3.95E-06 | -5.42E-06 | -8.01E-06 | -6.67E-06 | -3.40E-06 |  |
| 12 | 1.09E-06 | -1.35E-05 | -1.00E-06 | -2.13E-06 | -1.64E-05 | -2.15E-05 | -9.48E-06 | -2.46E-05 | -1.30E-05 | -6.45E-07 |  |
| 13 | 1.41E-06 | -1.19E-05 | -2.04E-06 | -3.78E-06 | -2.70E-06 | -4.94E-06 | -1.49E-05 | -1.66E-05 | -1.51E-06 | -5.09E-06 |  |
| 14 | 3.37E-07 | -6.08E-06 | -1.28E-06 | -4.77E-06 | -3.75E-06 | -4.77E-06 | -1.32E-05 | -9.62E-06 | -3.61E-07 | -3.97E-06 |  |
| 15 | 4.76E-08 | -1.27E-05 | -7.53E-07 | -1.03E-05 | -1.62E-05 | -4.54E-06 | -1.32E-05 | -8.29E-06 | -7.85E-06 | -1.85E-05 |  |
| 16 | 9.43E-07 | -2.52E-05 | -2.24E-06 | -2.01E-05 | -2.46E-05 | -8.93E-06 | -1.13E-05 | -7.21E-06 | -5.22E-06 | -1.00E-05 |  |
| 17 | 9.14E-07 | -1.81E-05 | -3.60E-06 | -9.74E-06 | -5.71E-06 | -8.71E-06 | -2.01E-05 | -1.35E-05 | -9.17E-06 | -4.23E-06 |  |
| 18 | -7.63E-07 | -7.49E-06 | -5.70E-06 | -7.17E-06 | -8.92E-06 | -6.24E-06 | -1.76E-05 | -7.27E-06 | -5.76E-06 | -3.54E-06 |  |
| 19 | 8.82E-07 | -4.91E-06 | -1.73E-06 | -6.77E-06 | -6.34E-06 | -1.36E-06 | -8.57E-06 | -8.63E-06 | -4.95E-06 | -2.52E-06 |  |
| 20 | 1.20E-06 | -1.97E-05 | -8.42E-07 | -4.42E-06 | -7.17E-06 | -1.26E-05 | -1.71E-05 | -6.13E-06 | -2.01E-06 | -5.36E-06 |  |
| 21 | 9.91E-07 | -5.45E-06 | -1.50E-05 | -8.07E-06 | -2.58E-06 | -7.09E-06 | -6.11E-06 | -9.21E-06 | -1.21E-05 | -6.17E-06 |  |
| 22 | 1.77E-07 | -1.17E-05 | -9.72E-06 | -9.78E-06 | -1.14E-05 | -1.80E-05 | -8.54E-06 | -1.66E-05 | -1.68E-05 | -4.43E-06 |  |
| 23 | 4.25E-07 | -8.23E-06 | -1.14E-05 | -1.09E-05 | -1.83E-05 | -4.16E-06 | -4.90E-06 | -4.27E-06 | -5.22E-06 |  |  |
| 24 | 1.04E-06 | -1.72E-05 | -7.18E-06 | -8.74E-06 | -1.34E-05 | -1.06E-05 | -4.31E-06 | -2.46E-06 | -2.48E-06 |  |  |
| 25 | 2.03E-06 | -7.96E-06 | -1.05E-05 | -1.06E-05 | -5.50E-06 | -5.84E-06 |  | -1.84E-06 | -3.11E-06 |  |  |
| 26 | 2.47E-06 | -2.03E-05 | -1.10E-05 | -4.15E-06 | -6.30E-06 | -1.37E-06 |  |  | -4.11E-06 |  |  |
| 27 | 1.67E-06 | -1.61E-05 | -2.47E-06 | -2.11E-06 | -5.16E-06 | -1.29E-05 |  |  | -3.71E-06 |  |  |
| 28 |  | -1.00E-05 | -2.04E-06 | -5.96E-06 | -6.53E-06 | -8.66E-06 |  |  | -6.75E-07 |  |  |
| 29 |  | -1.15E-05 | -4.18E-06 | -9.79E-06 | -6.55E-06 | -5.42E-06 |  |  | -5.62E-06 |  |  |
| 30 |  | -6.02E-07 | -3.38E-06 | -1.34E-05 | -8.19E-06 |  |  |  |  |  |  |
| 31 |  | -1.70E-05 | -5.02E-06 |  |  |  |  |  |  |  |  |
| 32 |  | -2.05E-05 | -3.67E-06 |  |  |  |  |  |  |  |  |
| 33 |  | -1.89E-05 | -9.18E-07 |  |  |  |  |  |  |  |  |
| 34 |  | -9.57E-06 | -3.62E-06 |  |  |  |  |  |  |  |  |
| 35 |  | -1.17E-05 | -3.43E-06 |  |  |  |  |  |  |  |  |
| 36 |  | -8.53E-06 | -1.24E-06 |  |  |  |  |  |  |  |  |
| 37 |  | -1.13E-05 | -2.71E-06 |  |  |  |  |  |  |  |  |
| 38 |  | -1.06E-05 | -4.93E-06 |  |  |  |  |  |  |  |  |
| 39 |  | -1.42E-05 | -5.49E-06 |  |  |  |  |  |  |  |  |
| 40 |  | -2.87E-05 | -2.68E-06 |  |  |  |  |  |  |  |  |
| 41 |  | -8.26E-06 | -3.19E-06 |  |  |  |  |  |  |  |  |
| 42 |  | -4.54E-06 | -5.26E-06 |  |  |  |  |  |  |  |  |
| 43 |  | -1.40E-05 | -1.16E-05 |  |  |  |  |  |  |  |  |
| 44 |  | -4.66E-06 | -1.69E-05 |  |  |  |  |  |  |  |  |
| 45 |  | -6.29E-06 | -4.52E-06 |  |  |  |  |  |  |  |  |
| 46 |  | -3.98E-06 | -2.55E-06 |  |  |  |  |  |  |  |  |
| 47 |  | -2.14E-05 | -9.78E-06 |  |  |  |  |  |  |  |  |
| 48 |  | -1.69E-05 | -1.08E-05 |  |  |  |  |  |  |  |  |
| 49 |  | -6.15E-06 | -7.42E-06 |  |  |  |  |  |  |  |  |
| 50 |  | -8.91E-06 | -6.04E-06 |  |  |  |  |  |  |  |  |
| 51 |  |  | -4.72E-06 |  |  |  |  |  |  |  |  |
| 52 |  |  | -2.52E-06 |  |  |  |  |  |  |  |  |
| 53 |  |  | -1.60E-05 |  |  |  |  |  |  |  |  |
| 54 |  |  | -6.10E-07 |  |  |  |  |  |  |  |  |
| 55 |  |  | -3.27E-07 |  |  |  |  |  |  |  |  |
| 56 |  |  | -1.34E-05 |  |  |  |  |  |  |  |  |
| 57 |  |  | -1.66E-06 |  |  |  |  |  |  |  |  |
| 58 |  |  | -4.36E-06 |  |  |  |  |  |  |  |  |
| 59 |  |  | -2.81E-06 |  |  |  |  |  |  |  |  |

*Experimental group brief descriptions:*

*Skeleton:* Coral skeleton, no living tissue, symbionts, or bacteria.

*Cryo-injured:* Negative control. Coral exposed to freezing in CVS1 solution via LN2 processing in an unsealed isochoric chamber. Evaluated 24h post-thaw / rehydration.

*Bleached Control:* Positive control. Healthy bleached (symbiont-free) coral.

*Tox3, Tox5, Tox7.5:* Corals exposed to standard CPA loading protocol, with final exposure time to full-strength CVS1 of 3, 5, or 7.5 minutes accordingly. Evaluated 24h after rehydration.

*Iso3, Iso5, Iso7.5:* Corals vitrified under isochoric conditions, after standard CPA loading protocol with final exposure time to full-strength CVS1 of 3, 5, or 7.5 minutes accordingly. Evaluated 24h post-thaw / rehydration.

**References**

1.	Daly, J. *et al.* Successful cryopreservation of coral larvae using vitrification and laser warming. *Sci. Rep.* **8**, (2018).

2.	Steif, P. S., Palastro, M. C. & Rabin, Y. The Effect of Temperature Gradients on Stress Development during Cryopreservation via Vitrification. *Cell Preserv. Technol.* **5**, 104–115 (2007).

3.	Steif, P. S. *et al.* Cryomacroscopy of Vitrification II: Experimental Observations and Analysis of Fracture Formation in Vitrified VS55 and DP6. *Cell Preserv. Technol.* **3**, 184–200 (2005).

4.	Sharma, A. *et al.* Cryopreservation of Whole Rat Livers by Vitrification and Nanowarming. *Ann. Biomed. Eng.* (2022). doi:10.1007/s10439-022-03064-2

5.	Gao, Z. *et al.* Vitrification and Rewarming of Magnetic Nanoparticle-Loaded Rat Hearts. *Adv. Mater. Technol.* **7**, 2100873 (2022).

6.	Couchman, P. R. & Karasz, F. E. A Classical Thermodynamic Discussion of the Effect of Composition on Glass-Transition Temperatures. *Macromolecules* **11**, 117–119 (1978).

7.	Fox, T. G. Influence of diluent and of copolymer composition on the glass temperature of a polymer system. *Bull. Am. Phs. Soc.* **1**, 123 (1952).

8.	Hallbrucker, A., Mayer, E. & Johari, G. P. Glass-liquid transition and the enthalpy of devitrification of annealed vapor-deposited amorphous solid water: a comparison with hyperquenched glassy water. *J. Phys. Chem.* **93**, 4986–4990 (1989).

9.	Plitz, J., Rabin, Y. & Walsh, J. R. The Effect of Thermal Expansion of Ingredients on the Cocktails VS55 and DP6. *Cell Preserv. Technol.* **2**, 215–226 (2004).

10.	Böhmer, R., Ngai, K. L., Angell, C. A. & Plazek, D. J. Nonexponential relaxations in strong and fragile glass formers. *J. Chem. Phys.* **99**, 4201–4209 (1993).

11.	Chen, T., Fowler, A. & Toner, M. Literature Review: Supplemented Phase Diagram of the Trehalose–Water Binary Mixture. *Cryobiology* **40**, 277–282 (2000).

12.	Rabin, Y. Mathematical modeling of surface deformation during vitrification. *Cryobiology* **102**, 34–41 (2021).

13.	Solanki, P. K. & Rabin, Y. Perspective: Temperature-Dependent Density And Thermal Expansion Of Cryoprotective Agents. *Cryoletters* **43**, 1–9 (2022).

14.	Tyree, T. J., Dan, R. & Thorne, R. E. Density and electron density of aqueous cryoprotectant solutions at cryogenic temperatures for optimized cryoprotection and diffraction contrast. *Acta Crystallogr. Sect. D Struct. Biol.* **74**, 471–479 (2018).

15.	Page, C. A., Muller, E. M. & Vaughan, D. E. Microfragmenting for the successful restoration of slow growing massive corals. *Ecol. Eng.* **123**, 86–94 (2018).

16.	Lager, C. *et al.* Metrics of Coral Microfragment Viability. *bioRxiv* (2023). doi:https://doi.org/10.1101/2023.01.03.522625
